# Lifestyle transition from rural to urban setting changes the gut bacterial profile in an ethnic community of northeast India

**DOI:** 10.1101/2025.07.11.664413

**Authors:** Santanu Das, Anupam Bhattacharya, Dibyayan Deb, Mojibur R. Khan

## Abstract

The human gut microbiome undergoes transformation due to various exogenous and endogenous influences. Lifestyle has been one of the major contributors of gut microbiome composition. Our study investigated the effect of migration on the gut microbiome of the *Mishing* ethnic community in India associated with urban migration from their pastural lifestyle. Gut bacterial diversity was profiled using 16S rRNA amplicon sequencing. We observed notable alterations in microbial diversity and composition, including changes in the core microbiome. While *Bifidobacterium, Lactococcus* and *Streptococcus* significantly increased (p < 0.01) among the migrant population there was depletion in bacteria such as *Succinivibrio* and *Prevotella*. This change in core microbiome was further reflected on the functional profile of the bacterial community harbored by the urban and rural dwellers. While the rural population was enriched in fatty acid biosynthesis pathways, amino acid and Vitamin B2 biosynthesis pathways were more prominent in the microbiome of the urban dwellers. We further observed increase in diversity and richness of the microbiome following migration. Co-analysis with ASVs from food samples consumed by the *Mishing* rural population revealed that 51 ASVs among the rural dwellers were from a widely consumed beverage, *Apong* which were completely absent among the migrants. Our analyses highlight the dietary preferences, particularly abstinence from specific foods and beverages, may modulate the alterations in gut microbial profiles. While the cross-sectional design and amplicon sequencing limited our ability to assess temporal and functional changes, our findings highlight a measurable impact of lifestyle and dietary transitions on the gut microbiome.

## Introduction

The human gastrointestinal tract (GIT) is home to trillions of microbes, collectively called ‘gut microbiome’[1,2]. In addition to digestion and nutrient assimilation, gut microbiome also plays important role in priming the immune system. The gut microbial composition within the GIT undergoes rapid, and in some cases, irreversible changes in response to dietary and environmental factors, physiological state and diseases (David et al., 2014; Ley et al., 2006; Wu et al., 2011b, 2011a). With the advent of modernization and industrialism, the human diet has undergone numerous changes from humble subsistence level food sources to highly processed food products [3]. In addition to diet and lifestyle, exposure to other factors associated with modernization may also significantly alter the gut microbial composition [7,8]. Many studies have compared the gut microbiome of urban and rural dwellers. In general, it is perceived that the urban dwellers have reduced microbial community and higher F/B ratio, and the rural population harbors a rich microbial diversity. [9]. Previous studies on comparison of urban and rural microbiome have focused mainly on the differences arising due to geographical location, and population undergoing transition. A study on Colombian population highlighted that transitioning population from rural to westernized has signatures of both hunter gatherers and industrialized countries [10]. Similarly, an Indian study on comparison of gut microbiome among rural and urban individuals reported the influence of topography and diet [11]. Although, previous studies have reported the change in gut microbiome due to migration, highlighting the effects of westernization, but reports on effect of migration within a single ethnic community is scanty . Therefore, to address this, we focused on investigating the effect of migration within a same community where genetic factors were constant. We studied the gut microbiome of a subpopulation of the *Mishing* community, who have migrated from rural areas to an urban and hence have undergone lifestyle transition and change in dietary pattern. Our previous study on the *Mishing* population, influence of a single dietary factor (alcoholic rice beverage) on the gut microbiome was observed [12]. This subpopulation of *Mishing* community had migrated from the northern bank of the river Brahmaputra (GPS Coordinates: 27.740790, 95.154990, 27.731543, 95.177729) in the state of Assam to Guwahati (GPS Coordinates: 26.128988, 91.827547) (state capital of Assam in Northeast India). due to recurrent flood and subsequent loss of agricultural land and habitats. Here we aim to identify the alterations in the gut bacterial profile among the migrants by comparing with their counterparts residing in rural area having agrarian lifestyle. To the best of our knowledge, our study is first of its kind which explores the lifestyle transition from rural to urban within a same ethnicity.

## Materials and methods

### Ethics statement

Our study was approved by the Human Ethics committee of the Institute of Advanced Study in Science and Technology (IASST) (Approval Number: IEC(HS)/IASST/1082/2014-15/6). Recruitment of volunteers was conducted adhering to the guidelines and protocols defined by the study. Additionally, all the participants were informed about the study both in verbal and written forms and their consents were taken prior to survey and sample collection.

### Study sites, volunteers, and sample collection

For our study, prospective volunteers were identified through the electoral database of Indian government. Based on the electoral data, a survey was conducted in areas dominated by *Mishing* community to initiate the volunteer recruitment and sample collection. The volunteers recruited were apparently healthy and belonged to the same ethnic community. The striking differences among the urban and rural dwellers were mainly reflected in their lifestyle and diet. The rural based community were primarily agrarian, and their diet depends on the seasonal crops. Primarily, the rural community consumed rice, seasonal vegetables, herbs, freshwater fishes, along with occasional consumption of meat (red meat and poultry) and pulses/lentils. The rural community also included fermented foods and beverages in their daily diet, primarily fermented fish (*Namsing*) and an artisanal alcoholic rice beverage called *Apong* (either *Poro* or *Nogin* based on their personal preferences). On the other hand, the urban dwellers earn their livelihood by working in industries, etc. Although the dietary pattern remains similar but relies on food products available in the local market. Interestingly, a few of the traditional food items such as *Apong* (alcoholic rice beverage) and *Namsing* (fermented fish) were in negligible quantities in the diet of the urban dwellers. In addition to difference in the lifestyle and diet, the urban dwellers are also exposed to pollutants and other factors associated with industrialization which affects the gut microbiome. The data for rural population were previously published in our previous study and can be accessed from NCBI-SRA under the accession number PRJNA906264.

A bi-lingual (*Assamese* and English) survey questionnaire was used to collect information from the volunteers. The questions were framed to gain insight into the dietary pattern, medical history, family lineages and demography. The inclusion criteria were *Mishing* ethnicity, age (18-50) and residence in the location for at least 5 years. The exclusion criteria include consumption of antibiotics and probiotics for the past 90 days, use of health supplements and based on their medical history.

Samples were self-collected by the volunteers. The volunteers were provided with a customized kit, which contained a sterile clinicols filled with 3ml RNA later for faecal sample collection, a pair of gloves and a disinfectant patch. The first defecation of the day was only considered as valid sample. After collections, the samples were immediately transported to the laboratory in frozen condition. On reaching the laboratory, samples were stored at -80 ℃ until processed. In addition to the samples, we have also collected anthropometric data such as height, weight, BMI, and blood groups during the sample collection. The demographic information of the volunteers is presented in the Supplementary Table 1. Samples and anthropometric data were collected under strict medical supervision.

### DNA extraction from faecal sample

DNA extraction was performed within 72 hours of sampling. Metagenomic DNA from the faecal samples were extracted using the Qiagen QIAamp PowerFecal Pro DNA Kit (Cat. No: 51804 Qiagen Inc, Germany) following the manufacture’s protocol. Briefly, the faecal samples were added to the bead beating tube for thorough homogenization. Following homogenization, the cells were lysed using mechanical and enzymatic treatment to release cellular contents. Genomic DNA was bound onto a silica column and purified using wash reagents supplied with the kit. Finally, genomic DNA was eluted in Tris EDTA buffer and stored at -80 ℃ until sequencing. Double stranded DNA (dsDNA) was quantified using One dsDNA kit with a Fluorometer (Quantiflour, Promega, Madison, USA).

### Amplicon library preparation and sequencing

The V3-V4 hypervariable region of the 16S rDNA was amplified using the primer pairs 341F-805R. For amplicon library preparation, amplification reaction consisted of 12.5 μl of 2× NEBnext Q5 hotstart mix (New England Biolabs, USA), 5 μl of 1pM forward and reverse primer pairs and 2.5μl of the template DNA. Amplification was carried out under the condition: 3 min at 95 °C (initial denaturation), followed by 25 cycles of 95 °C for 30s (denaturation), 55 °C for 30 s (annealing), and 72 °C for 30 s (extension) followed by a final extension at 72 °C for 5 min. Amplicons of desired size were selected using AMPure XP beads (Beckmann Coulter) in appropriate quantity. The purified amplicons were then subjected to index PCR that attaches dual indices and Illumina sequencing adapters using the Nextera XT Index Kit. The index PCR reaction consisted of a 50μl of reaction mixture containing 25μl of NEBnext Q5 hotstart mix (New England Biolabs, USA), 5μl of each of the Nextera XT Index 1 and 2 primers, 10μl of nuclease free water, and 5μl of template. Attachment of the indexes were performed under the following conditions: 3 min at 95°C (initial denaturation), followed by 8 cycles of 95°C for 30s (denaturation), 55°C for 30s (annealing), and 72°C for 30s (extension) followed by a final extension at 72°C for 5 min. Library of the desired size were validated using a capillary electrophoresis based fragment analyzer (Agilent Bioanalyser). Finally, the libraires were pooled in equimolar concentrations and sequenced on an in-house Illumina MiSeq instrument (Illumina Inc. USA).

### Bioinformatics analyses of the amplicon dataset

The paired end reads generated from Illumina sequencing were processed using the LotuS2 pipeline, with slight modifications of the default parameters [13]. Reads having less than 200 bases in length were discarded from downstream analyses. Sequences were clustered into Amplicon Sequence Variants (ASVs) using the DADA2 algorithm within Lotus2 [14]. Furter, sequence clusters were refined for human contamination and crosstalk using LULU and UNCROSS2, respectively [15,16]. ASVs obtained were aligned with Lambda against the human intestinal (HITdb) and SILVA 138.1 databases [17,18]. For removal of the sequencing primers -forwardPrimer and -reversePrimer options in the LotuS2 pipeline was used. Out of 58,428,184 reads 48,840,843 reads were classified as “High-quality”. A total of 9,587,326 reads discarded as they did not conform to the criteria (Q ≥ 30 and read length = 200 bp). The good quality reads generated a total of 6437 ASVs, which were retained in the matrix for diversity and taxonomic analysis.

Downstream analyses of the amplicon data were carried in R (v 4.4.1) environment using the phyloseq and microeco pacakages [19,20]. For calculation and estimation of diversity index vegan package (v 2.6-8) was used [21]. Functional potential of the bacterial community was profiled using Picrust2 pipeline [22].

### Statistical analyses

All the statistical tests were performed within the R environment (v 4.4.1) utilizing the base functions. Specialized packages such as vegan, microeco, microbiome and microbiomeutilities [19,21,23] were used for diversity analysis and visualization. For comparisons of the microbial diversity, the samples were rarefied to an equal depth (22011). Thereafter, the diversity and richness (alpha diversity) were calculated for each of the samples which includes Chao1, number of observed features and Shannon indices. Prior to computing the beta diversity, log transformation was performed for consistency. We used the “adonis2” function in the vegan package to determine the statistical significance. Further, “betadisper” function in the vegan package was used to estimate the homogeneity. For determining the statistical significance, we used Kruskal-Wallis test with Bonferonni-Hoschberg (BH) corrections. The differential abundance of taxa between the rural and urban population was determined through Aldex2 and LEfSe methods [24,25]. For differential analysis, we considered p-value of 0.05 after false discovery rate correction utilizing the BH method.

## Results

### Lifestyle transition from rural to urban has significant effect on gut microbiome composition

We accessed the bacterial diversity of both the rural and urban dwellers. It was observed that three major phyla dominated the bacterial composition (> 90 %) irrespective of the habitats (Figure 1 A). However, the composition of the phyla varied considerably among the groups. Bacteroidetes which was 42.40% among the rural dwellers was only 30.01% among the urban residents (p > 0.0001). This void in Bacteroidetes was compensated by abundance of Firmicutes, which had occurred at an abundance of 48.67% as opposed to 28.53% in the rural population (p > 0.0001) (Supplementary Figure 1A and 1B). Interestingly, the phyla Proteobacteria which contains many pathogens and pathobionts was abundant in rural population (16.96%) as opposed to urban dwellers (5.61%) (p < 0.001), raising a serious concern about the sanitization condition prevalent in the rural areas. Interestingly, we observed that the phyla Actinobacteria, was significantly higher among the urban dwellers (3.73%) as compared to rural population (0.089 %) (Figure 1B). Minor phyla such as Elusimicrobio, Tenericutes, Spirochates, and Verrucomicrobia were present both in rural and urban population (Figure 1A).

**Figure 1.**
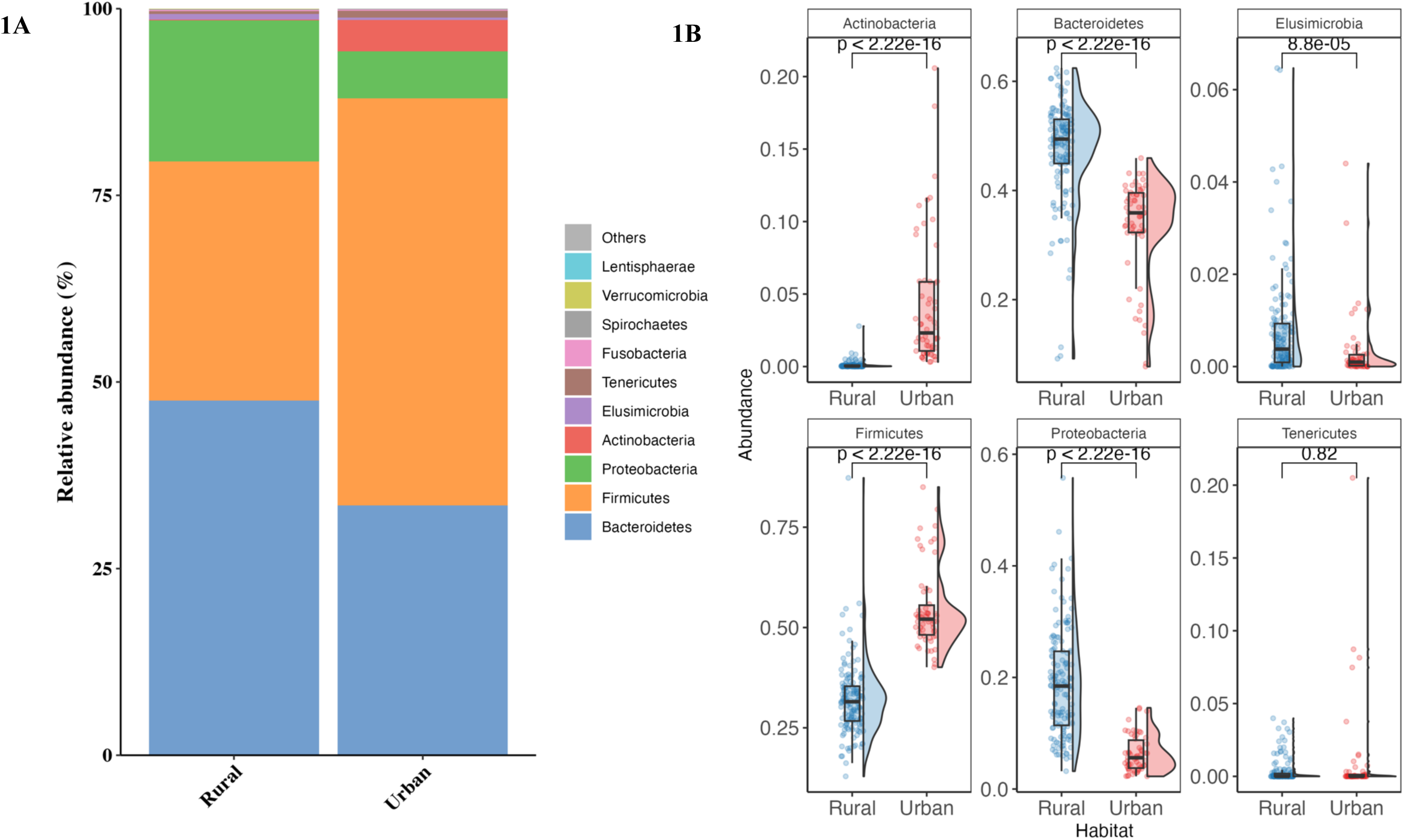
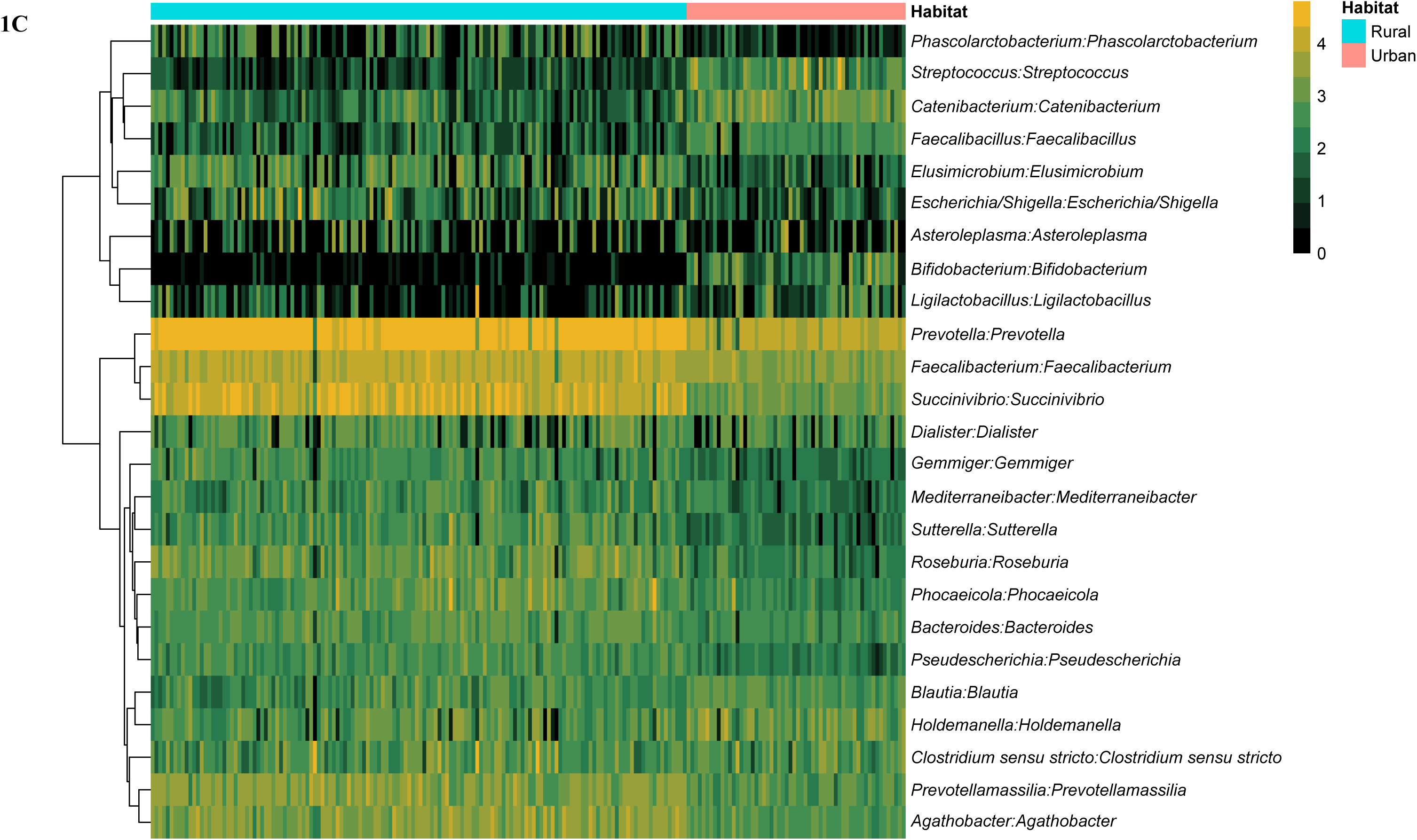
Microbiota composition of the rural and urban dwellers. (A) Relative abundance of the top 10 dominant phyla. (B) Differential abundance of the major phyla between the rural and the urban population. (C) Heatmap demonstrating the most abundant genera across the rural and urban populations. The relative abundance of the genera is scaled from 0 to 4 represented by green to yellow colour, where green signifies lower and yellow signifies higher abundances.

Further resolution of the taxonomic composition to genus level revealed that irrespective of living conditions, the genus *Prevotella* was the most dominant bacteria, which is widely present among any Indian population. Consistent with the observations made at the phylum level, the top 10 prominent genera varied in abundance, among rural and urban population. *Prevotella*, which was the most prominent bacteria, was more abundant among rural population (28.53%) as compared among the urban dwellers (17.49%). Our previous studies have reported the dominance of *Succinivibrio* among the ethnic communities in Assam. In this study we observed that the migration to urban areas results in significant reduction in the abundance of *Sucinivibrio*. (12.67% to 3.09%) (Figure 1C). While *Faecalibacterium*, *Agathobacterium*, *Prevotellamassila* and *Roseburia* were more abundant in rural population; *Ligilactobacillus, Blautia, Bifidobacterium and Holdemenella* were more abundant among the urban populations (p < 0.05).

### Urban lifestyle enriches the abundance of Firmicutes and Actinobacteria

For deeper insight into the gut bacterial differences between the rural and urban population, we extended the differential abundance test to the ASV level. It was observed that a total of 1687 ASVs were differentially abundant (adjusted p-value < 0.05), with a larger number of ASVs enriched among the urban dwellers (Figure 2A). Majority of the ASV enriched in the urban population belonged to the Firmicutes, followed by Bacteroidetes and Proteobacteria. Additionally, we also assessed the effect size of these differentially abundant ASVs to determine the magnitude of their contribution to microbiome variation between two groups. Effect size analysis revelated that 43 amongst the significantly different ASVs had a fold change greater than 2 (Log_2_Fold Change > 1) (Figure 2B), indicating substantial shift in abundance. Effect size analysis further revealed that highly significant ASVs (adjusted p-value < 0.01) also exhibited large effect sizes, reinforcing their biological relevance in differentiating the two populations (Supplementary Table 2).

**Figure 2.**
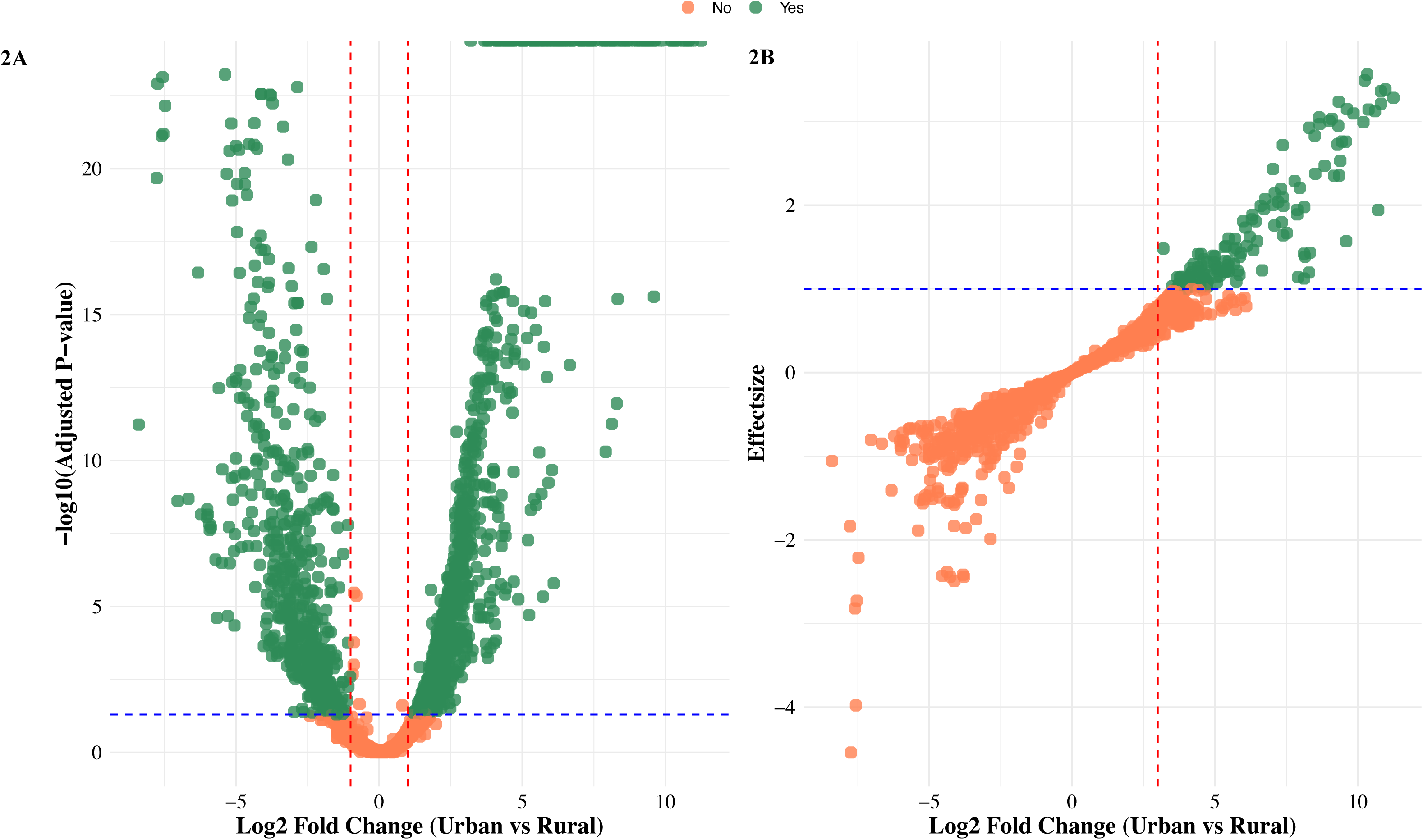
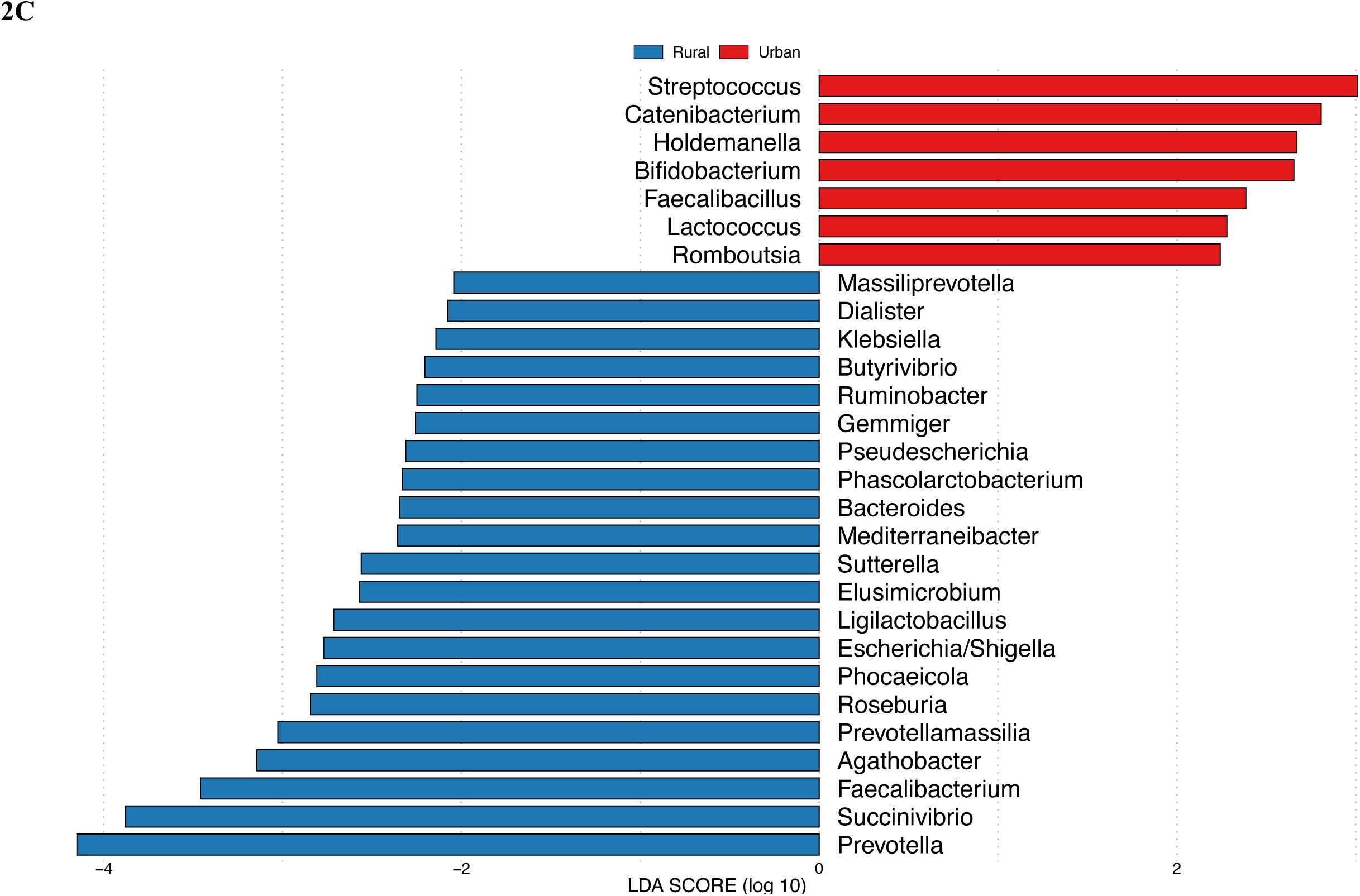
A) Volcano plot depicting the differentially abundant taxa in urban and rural population. X-axis represents the Log2 Fold Change (Urban vs. Rural) and the Y-axis represents the -log10(Adjusted P-value). ASVs with a positive Log₂ Fold Change indicate higher abundance in the Urban group, whereas those with a negative Log₂ Fold Change are more abundant in the Rural group. Non-significant ASVs are displayed in gray, while significantly different ASVs are marked in red. B) Effect size plot which indicate the magnitude of group differences, providing an estimate of the strength of association. ASVs with effect size more than 1 and Log2 Fold Change more than 2 are represented in green. C) Linear Discriminant Analysis (LDA) Effect Size (LEfSe) plot of signature taxa in rural and urban population. Taxa marked in blue are overrepresented among the rural dwellers, whereas taxa in red are overrepresented among the urban population

Next, we checked the differential abundance using Linear discriminant analysis Effect Size (LEfSe) to undermine the statistically significant biologically relevant signatures (genus) associated with each of the urban and rural lifestyle gradient. We observed that more taxa were enriched among the rural dwellers (n = 21) as compared to the urban dwellers (n = 7). Notably, *Succcinivibrio*, *Prevotella*, *Faecalibacterium*, *Agathobacterium*, and *Prevotellamassila* were highly enriched among the rural population (LDA score < 3). On the contrary, *Streptococcus*, *Catenibacterium*, *Holdemanella*, *Bifidobacterium*, *Faecalibacillus*, *Lactococcus, and Romboustia* were enriched among the urban dwellers (LDA score < 3) (Figure 2C). Our findings suggests that change in lifestyle from rural to urban may have impacted on their gut bacterial composition with implications for host metabolic functions.

### Gut bacterial community among the rural dwellers are more resilient

The core bacterial community among the rural and urban population was determined considering both prevalence and abundance. “Core microbiota/ core community” refers to the conserved taxa that is present in a majority of the individual in a community. Since, there is no defined criterion for determining the core, we considered a prevalence of 90 % and abundance of 0.01 for determining the “core” in our study. We observed marked difference in the core bacteria in rural and urban populations. While 13 genera made up the core bacteria in the urban community, the core bacteria in amongst the rural dwellers was more resilient and comprised of 9 genera only. These 9 genera include *Prevotella*, *Succinivibrio*, *Facecalibacterium*, *Agathobacter*, *Prevotellamassilla*, *Roseburia*, *Phocaeicola*, *Clostridium* and *Holdemenalla* (Figure 3A). However, among the core bacteria in the urban population, we observed taxa such as *Catenibacterium*, *Bacteroides, Blautia and Romboustia* (Figure 3B).

**Figure 3.**
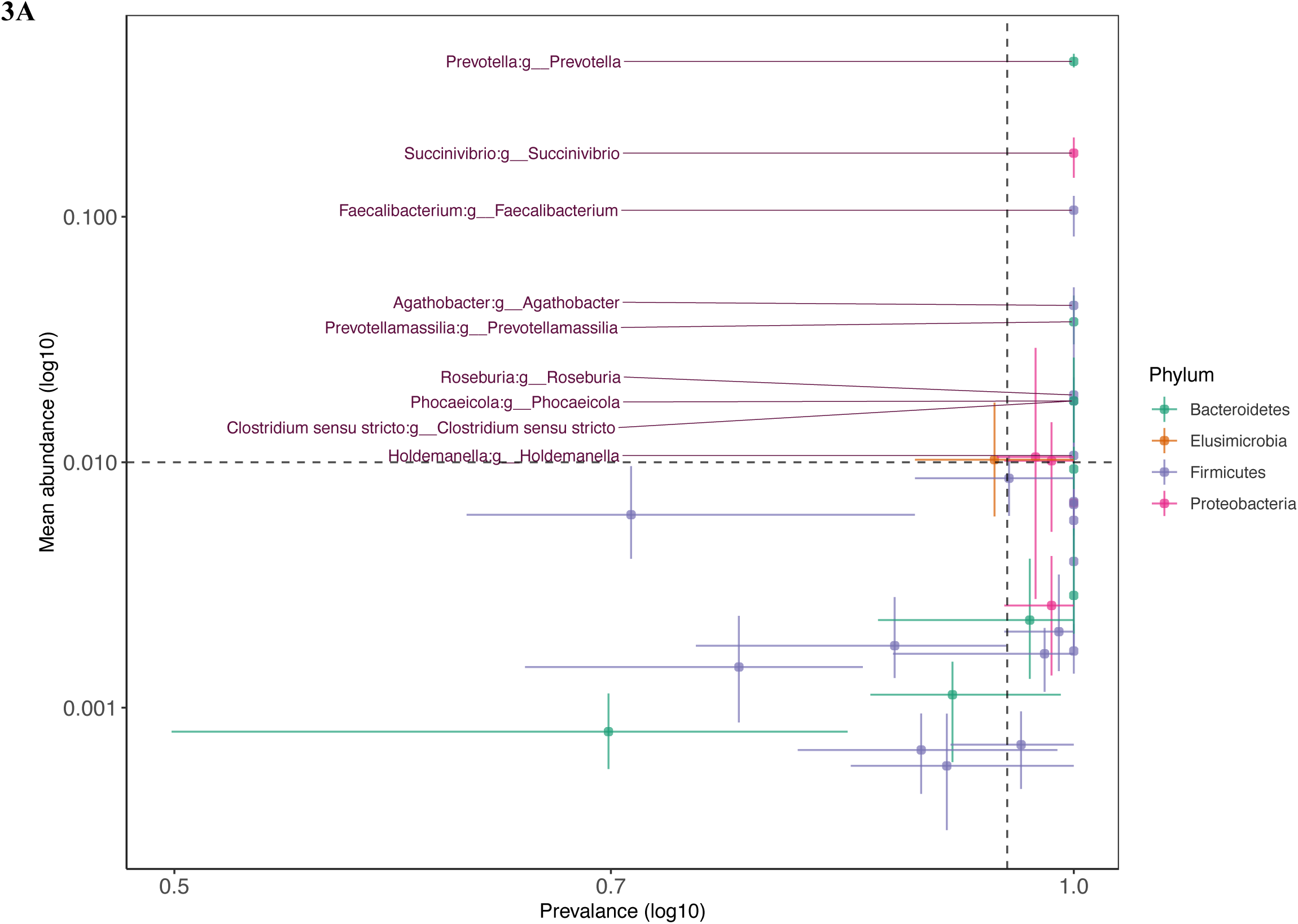
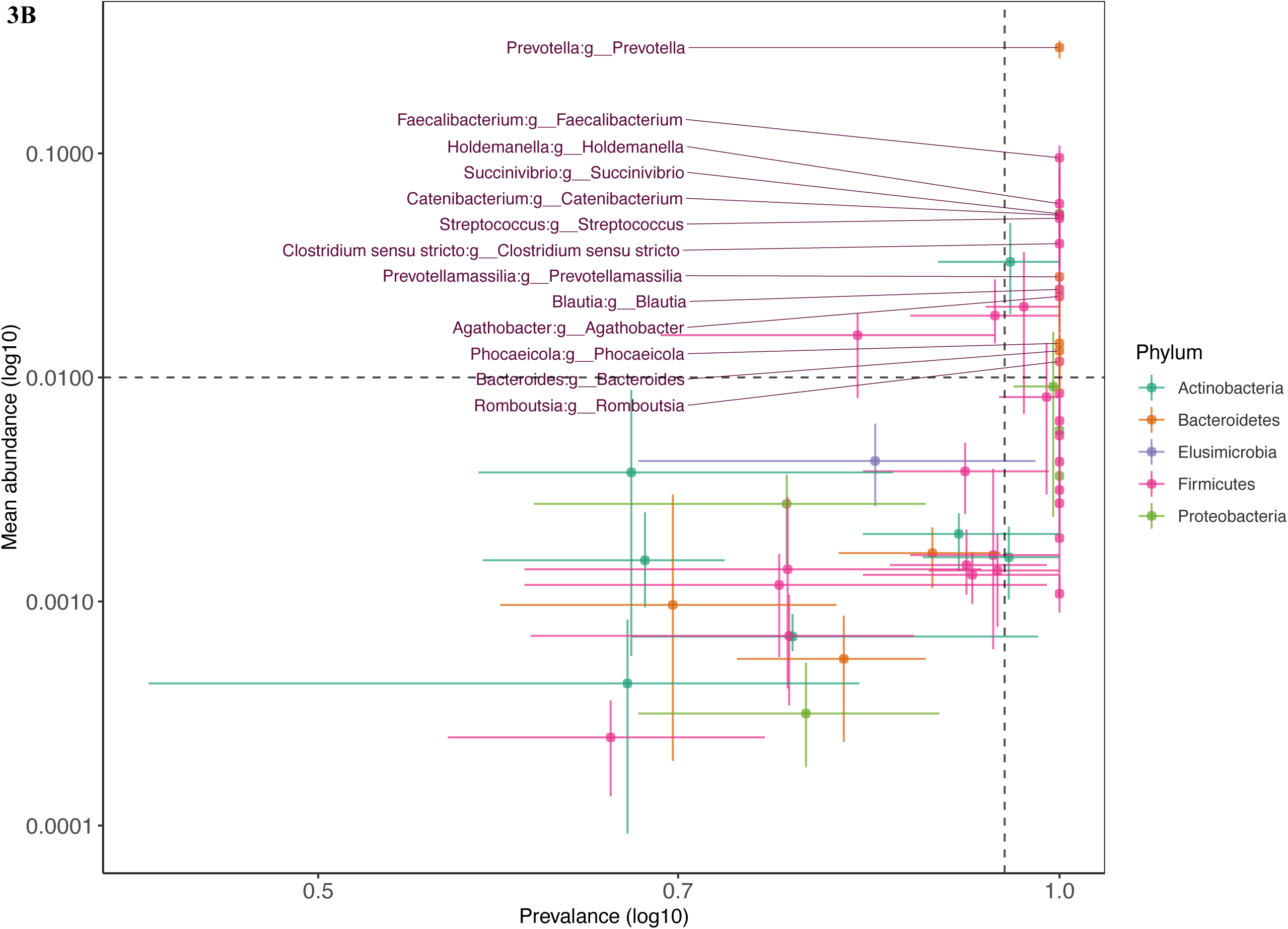
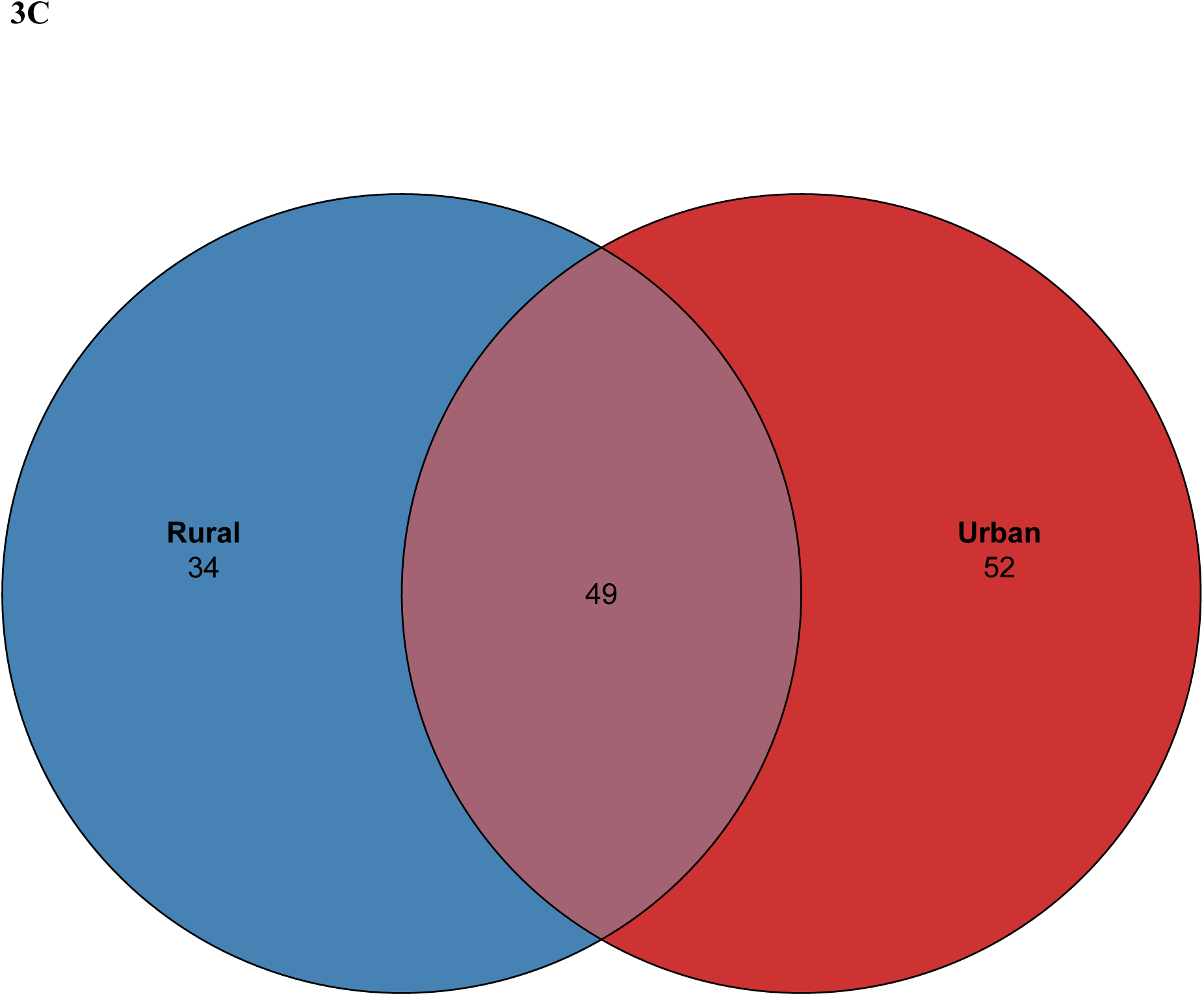
Core and shared microbiota A) Plot depicting the log10 prevalence (X-axis) and log10 abundance (Y-axis) among the rural population. Taxa with a log10 prevalence threshold > 0.9 and log10 abundance > 0.01 qualifies as core bacteria. B) Plot depicting the log10 prevalence (X-axis) and log10 abundance (Y-axis) among the urban population. Taxa with a log10 prevalence threshold > 0.9 and log10 abundance > 0.01 qualifies as core bacteria. C) Venn diagram depicting the abundance of shared and unique in the population. The intersection represents the number of shared taxa between urban and rural population.

Further we also investigated the unique and shared ASVs among the two group of habitation. Among the core ASVs, 49 ASVs were identified to be shared between the group of lifestyle gradient. These 49 ASVs were assigned to *Prevotella*, *Faecalibacterium*, and *Agathobacter*. However, we made an interesting observation that, transition to urban lifestyle drastically increases the number of unique ASV in the gut bacteria, as evident from 34 unique ASVs in urban to 52 unique ASVs in rural population (Figure 3C). The unique ASVs among the rural population were more assigned to *Blautia, Elusimicrobium*, *Gemmiger*, and *Succinivibrio*. Contrary to that the unique ASVs in the urban population were assigned to genera such as *Catenibacter, Clostridium, Dialister, and Phocaeicola*.

### Migration to urban lifestyle increases the richness and diversity of gut microbiota

For estimation of bacterial diversity, samples were rarefied to an even depth of 20,000 reads per samples (Supplementary Figure 2) and the bacterial diversities were estimated using three different diversity metrices to gain insight onto the richness and diversity. It was observed that transition from rural to urban lifestyle resulted in significant increase in overall microbial richness (Chao1 and Observed Features) and diversity (Shannon diversity) (Figure 4A,B,C). This finding concords with the findings of “core bacteria” where we observed resilience in the taxa among rural dwellers.

**Figure 4.**
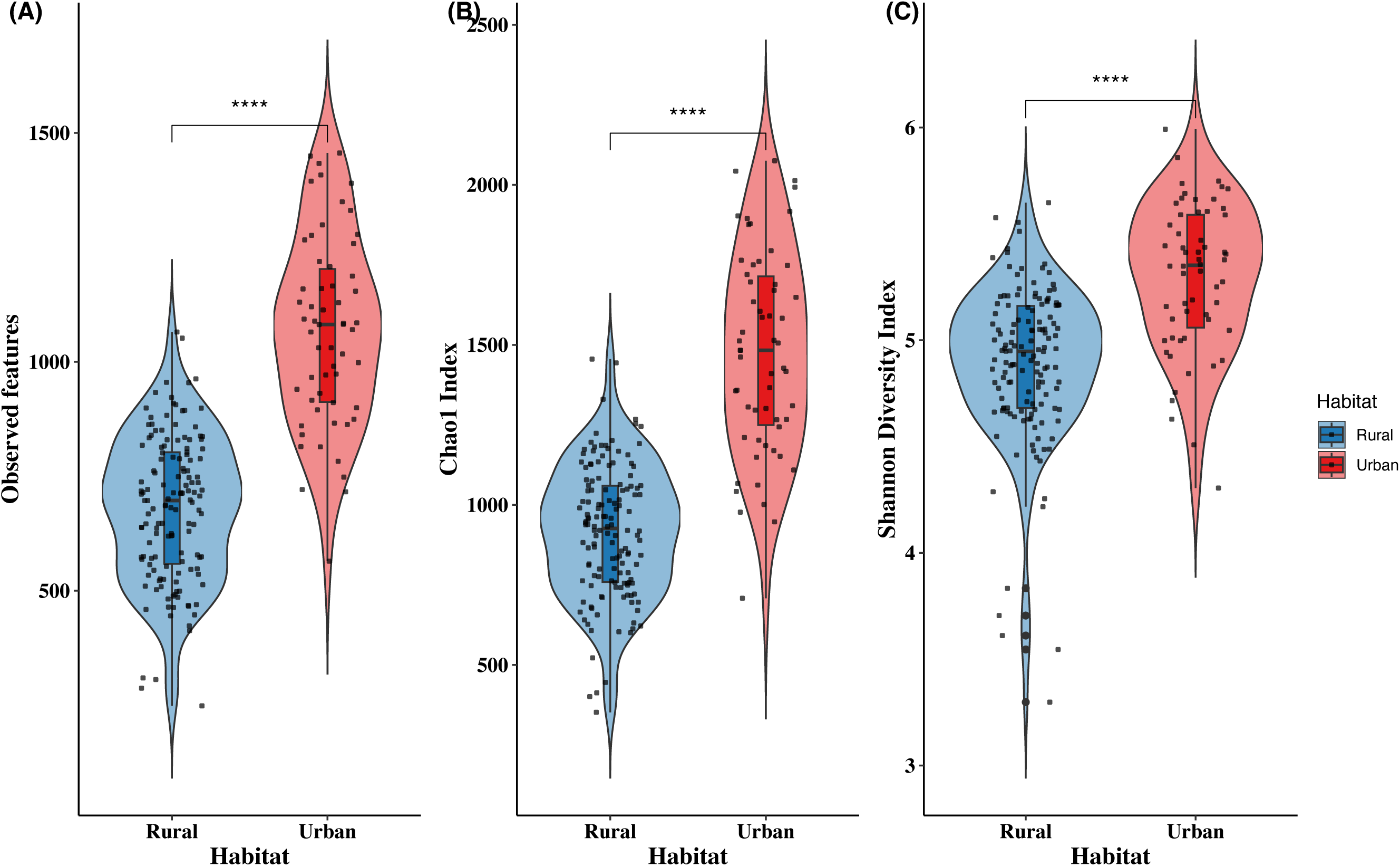
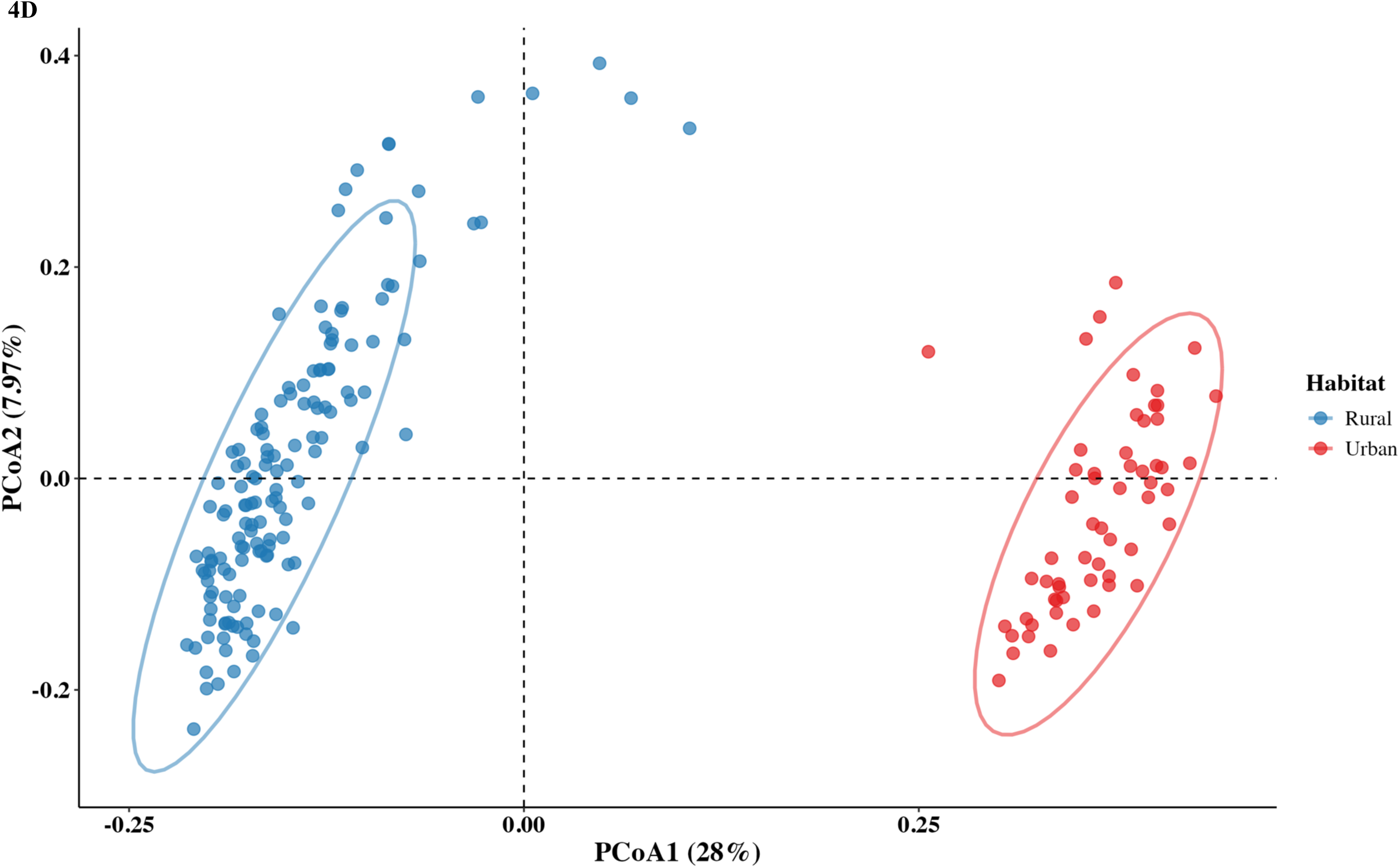
Microbial diversity (richness and diversity) as estimated by A) Observed features B) Chao1 index and C) Shannon diversity index. D) Principal coordinate analysis (PCoA) of the Bray-Curtis distances of the gut bacterial composition of rural and urban dwellers.

The compositional bacterial diversity was investigated by computing Bray Curtis and Weighted UniFrac distances. We observed prominent significant alternation in the composition of microbial diversities upon transition of lifestyles. In the PCoA plot, two distinct clusters were observed for rural and the urban population, explained by a variance of 28% and 7.97% for Axis1 and Axis2, respectively (Fig 4D). Interestingly, despite of having variation in the bacterial composition, we did not observe variance in the dispersion of microbial community among urban and rural dwellers. (Supplementary Figure 3).

### Lifestyle transition alters the functional potential of the gut bacterial community

To understand the functional potential of the bacterial community harbored by the rural and urban dwellers, we predicted the functional potential using PICRUSt2 tool. Interestingly, we observed remarkable difference in the functional potential of the bacterial communities in rural and urban population. In the PCoA plot based on the Bray Curtis distance of pathways, we observed distinct clusters for the urban and the rural population, separated by 65.3% and 11.1% for axes PCoA1 and PCoA2, respectively (PERMANOVA R^2^ = 0.5985, p =0.001). Further analysis of variance using beta dispersion revealed that the variation between the functional potential of the bacterial community among rural and urban dwellers was significantly larger with the communities having different dispersions (F = 13.809, p = 0.002) (Figure 5A). A total of 392 functional pathways (MetaCyc Metabolic Pathway Database) were identified in the bacterial community of both rural and urban dwellers. The top 50 functional pathways common to both the urban and rural bacterial community is represented in a heatmap (Figure 5B). The prominent functional pathways includes pathways for glucose utilization (ANAGLYCOLYSIS-PWY, CALVIN-PWY, NONOXIPENT-PWY and PWY-6737), glycogen biosynthesis and degradation (GLYCOCAT-PWY, GLYCOGENSYNTH-PWY), unsaturated fatty acid biosynthesis pathways (FASYN-ELONG-PWY, PWY-7663), amino acid biosynthesis pathways (ARO-PWY, BRANCHED-CHAIN-AA-SYN-PWY, ILEUSYN-PWY, PWY-2942, PWY-3001, PWY-5097, PWY-5101, PWY-5103, PWY5014, VALSYN-PWY). However, most of the common pathways are involved in cellular processes of the bacterial cell.

**Figure 5.**
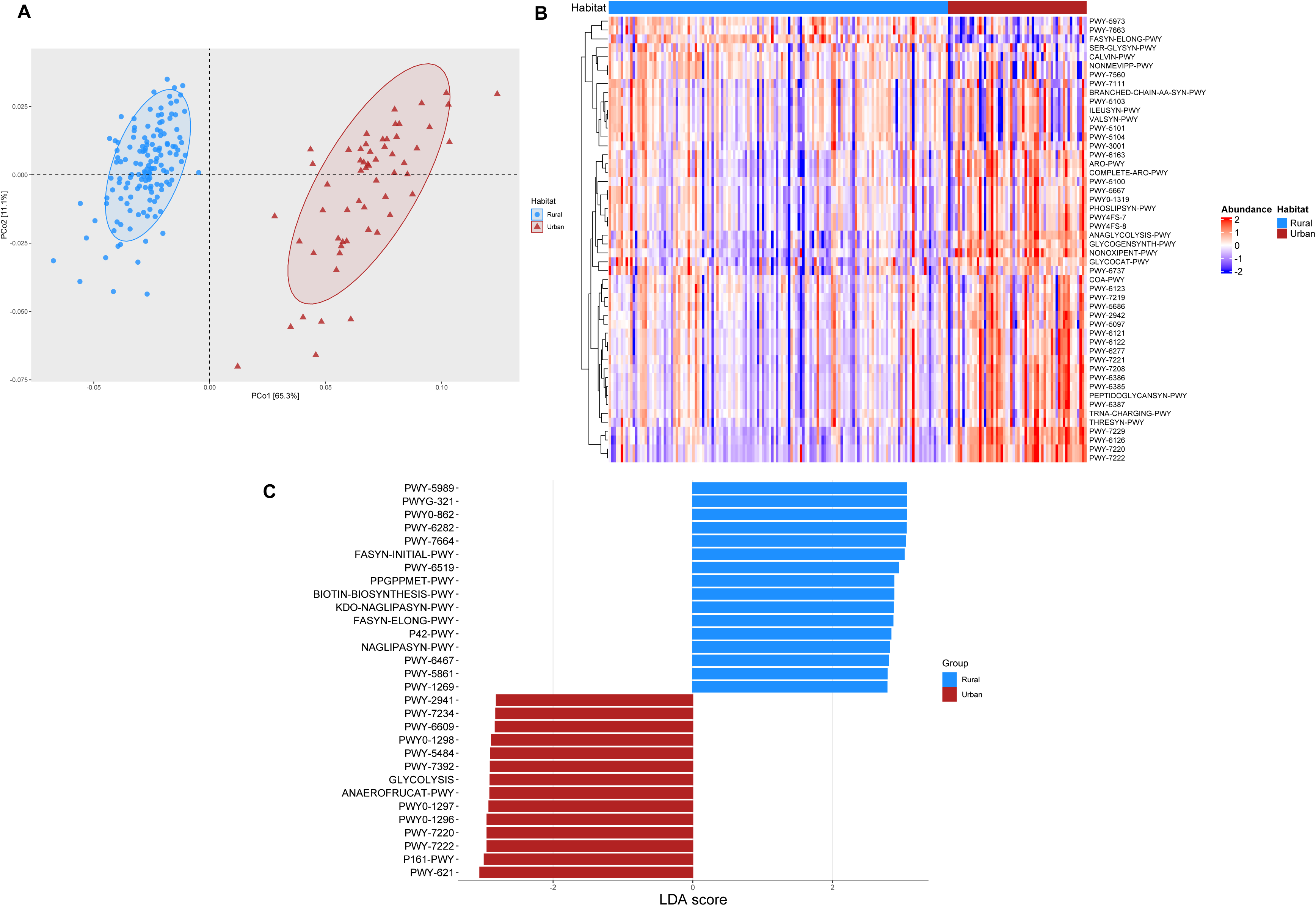
Predicted functional potential of the bacterial communities. (A) Principal coordinate analysis (PCoA) plot based on the Bray-Curtis distance of the functional pathways. The ellipses represent a 95% confidence interval. (B) Heatmap depicting the top 50 functional pathways in rural and urban population. The abundances of the pathways (Z-score) are scaled from -2 to +2 with blue colour representing lower abundance and red colour representing higher abundance. (C) Linear discriminate analyses (Lefse) of the differentially abundant pathways in rural and urban dwellers. The top 30 differentially abundant pathways are represented (LDA score (3.05-2.7).

Finally, to explore the difference in functional potential of the bacterial communities in rural and urban dwellers, we conducted Linear discriminant analysis Effect Size (LEfSe) (Figure 5C). We observed that out of the 392 pathways, 281 pathways were differentially abundant (LDA score 0.05-3.06, adjusted p-value < 0.01) (Supplementary Table 3). Pathways pertaining to fatty acid biosynthesis (PWY-5989, PWY0-862, PWY-6282 , PWY-7664, FASYN-INITIAL-PWY and FASYN-ELONG-PWY) were prominent in the bacterial community harbored by the rural population. Additionally, amino acid biosynthetic pathways for tyrosine, methionine, arginine, lysine and histidine (PWY-6630, HSERMETANA-PWY, HOMOSER-METSYN-PWY, PWY-5347, ARGSYNBSUB-PWY) and vitamin B2 biosynthesis pathway (RIBOSYN2-PWY, PWY-6167) were significantly enriched in the urban population. Contrary to that, metabolic pathways such as sucrose degradation (PWY-621), homolactic fermentation (ANAEROFRUCAT-PWY), glycolysis (PWY-5484), glycogen biosynthesis (GLYCOGENSYNTH-PWY), heterotactic fermentation (P122-PWY) and mixed acid fermentation (FERMENTATION-PWY) were more enriched among the urban dwellers. Interestingly, aspartate super pathway (PWY0-781) and L-lysine biosynthesis (PWY-2941) pathways were also enriched among the urban dwellers.

### Change in dietary pattern due to migration results in loss of certain bacteria in the gut microbiome

We traced back the difference of the bacteria communities to dietary habits followed by the *Mishing* community. In our previous studies we have reported that consumption of a fermented beverage (*Apong)* alters the gut bacteria within the same community. In this study, comparing the ASV of *Apong* to the ASV of gut bacteria using multiple sequence alignment tools, we identified the ASVs that might have probably been inherited from the drink consumed. For doing this, we created a database using the representative sequences of bacteria present in *Apong*. While none of the ASVs from the urban population matched with the ASVs present in *Apong,* we identified 51 ASVs from the rural population exactly matched with ASVs present in *Apong* (Supplementary Table 4). The majority of these ASV (n = 12) were assigned to *Succinivibrio* explaining the abundance of this bacterium in the gut microbiome of the rural *Mishing* population. Next, we also identified 10 of the shared ASVs belonging to Lactic Acid Bacteria (LAB) including *Lactobacillus, Pediocccus* and *Leuconostoc* which were prominent bacteria present in *Apong.* The rest of the matched ASVs were assigned to *Prevotella* and members of Enterobacteriaceae family.

## Discussion

The human gut microbiome is influenced by an array of factors such as birth mode, lifestyle, disease conditions ageing, etc of which lifestyle and diet are the most prominent ones [3]. In general, it is perceived that migration is associated with alternation with microbiome with potential change in functional attributes [26]. Additionally, studies comparing the gut microbiome among rural and urban dwellers have previously reported the enrichments of Proteobacterial genus among urban populations [11]. Our study focuses on unravelling the changes in gut microbiota of a population within the same ethnicity which have transitioned from rural to urban lifestyle with eventual loss of traditional dietary habits.

We found that the bacterial diversity differs demonstrably in lifestyle transition from rural to urban. The urban dwellers exhibited a more diverse gut microbial community compared to the rural dwellers, which had a stable community with fewer observed ASV and Choa1 diversity index. Additionally, we also found that within the “core”, 52 were exclusive only to the urban dwellers as opposed to 34 among the rural population. The urban specific ASVs belonged to *Mediterraneibacter* and *Catenibacterium* among others. *Mediterraneibacter* is usually associated with obesity and might be a signature of urban lifestyle [27]. *Catenibacterium* can ferment glucose into multiple short chain fatty acids [28]. Notably the shared bacteria between both the groups includes *Prevotella,* as the dominant bacteria, consistent with its prevalence among Indians [11,29]. The abundance in *Prevotella* can be attributed to frequent consumption of carbohydrates and green vegetables [28,30]. Other common bacteria in both the groups includes *Faecalibacterium*, *Succinivibrio, Agathobacter, Phocaeicola* and *Clostridium*. The dominance of both *Faecalibacterium* and *Agathobacter* can be attributed towards the fiber rich diet of *Mishing* community [31]. Despite being a core bacterium among the urban population, *Agathobacter* was highly elevated among the rural populations being a signature taxon in LEfSe analysis. The higher abundance of *Agathobacter* in rural populations is due to their frequent intake of plant-based fibers, which are often lacking in urban diets. Within the gut microbiota, *Agathobacter* indulges in cross feeding mechanisms to synthesize butyrate from acetate produced by other gut bacteria [31–33]. Similarly, *Phocaeicola* which have been recently re-classified from *Bacteroides* may produce succinate from complex carbohydrates which are otherwise undigested by human digestive system [34,35]. Initially *Phocaeicola* was suspected to be a pathogenic having role in multiple human infections [36]. However, recently it has been shown that 3-Hydroxyphenylacetic acid produced by this bacterium downregulates enzymes that promotes lipid storage in hepatocytes and thereby prevents metabolic disorders [37].

Our findings on the core bacteria highlights that transition in lifestyle have impacted the core bacteria among the migrants. However, certain bacteria such as *Prevotella*, *Succinivibrio*, *Agothobacter* etc. sustained to thrive. We speculate that, these bacteria maintain the ecology and functionality of the gut ecosystem. For example, genera such as *Succinivibrio* and *Phocaeicola* produce succinate from the complex carbohydrates in both the population; but those are utilized by different groups of bacteria such as *Roseburia* in rural and by *Catenibacterium* among the urban population to produce butyrate and other essential metabolites [28,38–40].

The differences in the core bacteria among the rural and the urban dwellers were also reflected in the functional potential exhibited by these bacterial communities. Enrichment in unsaturated fatty acid biosynthetic pathways for palmitoleate, enoate and oleate highlights the potential role of the bacterial community in synthesis of good fat with potential benefits of insulin sensitivity, lowering cholesterol and blood pressure [41–43]. Similarly, enrichment in pathways for biosynthesis of essential amino acids and vitamin shows the crucial role played by gut microbes in host nutrition and overall health [44,45]. Nevertheless, our analysis on the functional potential of the bacterial communities were based on reference genomes and phylogenetic placement to predict functional pathways and hence may not depict the actual functional roles of the bacterial community.

We observed that the genera *Bifidobacterium* and *Lactococcus* were highly abundant among the urban dwellers as compared to the rural population. While conducting our surveys, and sample collection we learned that milk and dairy products are scarcely consumed by the rural *Mishing* community. Contrary to this, the urban population included milk and dairy products in their dietary regime which might have enriched such bacteria. The enrichment of *Ligilactobacillus* among the rural population can be attributed to the frequent consumption of *Apong,* enriched with LABs .

Relating dietary habits to the bacterial abundance in the volunteers, we were able to trace back few of the ASVs to the fermented beverage *Apong*, consumed frequently by the rural population. Our previous studies have highlighted the abundances of LAB in *Apong* and the impact of frequent *Apong* consumption on gut health. In the current study, we observed that a few of the ASVs belonging to LAB were exact match of the ASVs present in *Apong.* Therefore, we assume that these bacteria were successful in establishing in the intestine of the rural population and have conferred their benefits. However, upon change in dietary pattern due to migration, these bacteria have eventually diminished from the intestine of the migrants, owing to the transient nature of LABs [46]. On the contrary, the similarity of ASVs of *Prevotella, Succinivibrio* and members of the Enterobacteriaceae family to that of *Apong* raises the concern of hygiene and sanitization prevalent in rural areas. This could be due to their use of untreated drinking water in the preparation of *Apong*.

## Conclusion

Our study presents the changes in human gut bacterial profile due to transition of lifestyle from rural to urban environment within a same ethnic community. Our observations revealed that migration leads to change in diversity and composition in the gut microbiome, with significant variation in the core microbiome composition. Further, correlation of the gut microbiome to the dietary factors revealed that abstinence from certain food and beverages fosters change within the gut microbiome. Taken together our finding suggests that habitation changes from rural to urban introduces subtle but significant changes in the gut microbiome. Being a cross-sectional study, we were unable to monitor the changes in microbiome over time. Moreover, we used 16S rRNA amplicon data to profile the diversity and hence were unable to explore the functional differences in the microbiota. A shot gun metagenomic DNA sequencing would have enabled us to establish a more causal relation on the transition of microbiome and their relation to dietary practice. Nevertheless, our study was successful in highlighting the impact of lifestyle changes on the gut microbiome within same ethnicity. To the best of our knowledge, our study is the first in Indian context to evaluate the effect of lifestyle transition on gut microbiome within the same community.

## Supplementary materials

All the supplementary files can be accessed from the file supplementary documents.pdf

## Data availability

Sequencing data are available on the NCBI SRA server under the BioProject ID PRJNA906264. The Biosample accession of the individual samples can be found on Supplementary Table 5. This study used sequencing samples from previously published studies, which can also be found under BioProject ID PRJNA906264.

## Author’s contribution

Santanu Das: Conceptualization, Formal analysis, Investigation, Methodology, Visualization, Writing-original draft; Anupam Bhattacharya: Formal analysis, Methodology, Dibyayan Deb: Methodology, Mojibur R. Khan: Conceptualization, Funding acquisition, Resources, Supervision, Writing – review & editing.

## Acknowledgements

The authors acknowledge the Advanced Level Institutional Biotech Hub of IASST and the Central instrumentation facility of IASST for providing the facilities. The authors also thank Mr. Basanta Mili for his assistance in recruitment of volunteers.

