## Supplementary document for "Lifestyle transition from rural to urban setting changes the gut bacterial profile in an ethnic community of northeast India"

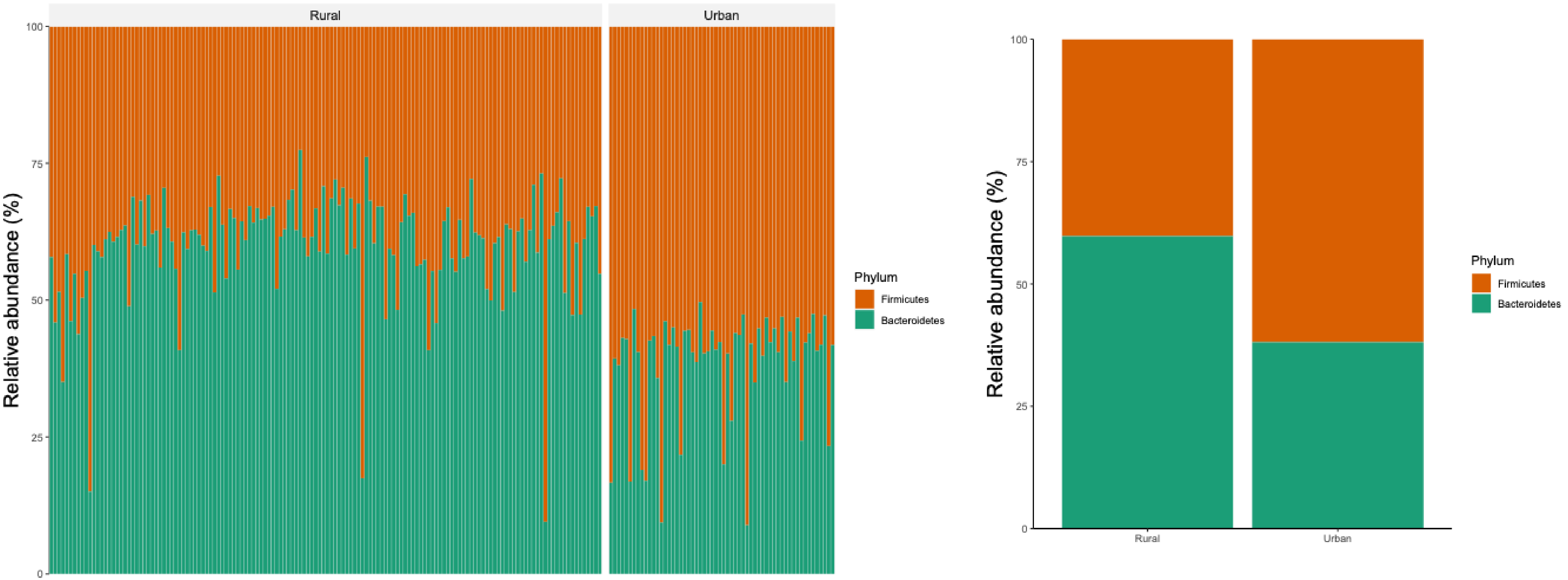


Supplementary Figure 1 : F/B ratio among the rural and urban dwellers (A) F/B ratio in across all samples faceted by habitat (B) Mean F/B ratios among rural and urban dwellers

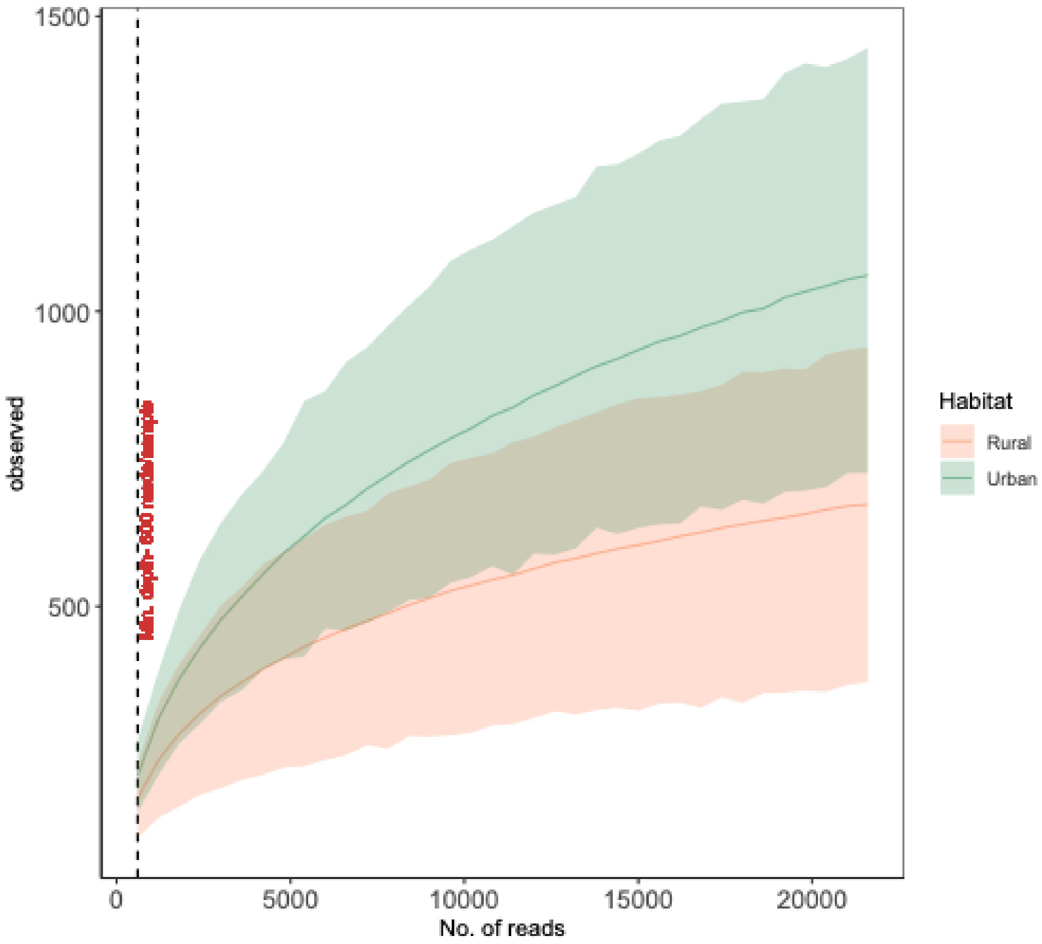


Supplementary Figure 2: Rarefaction curve

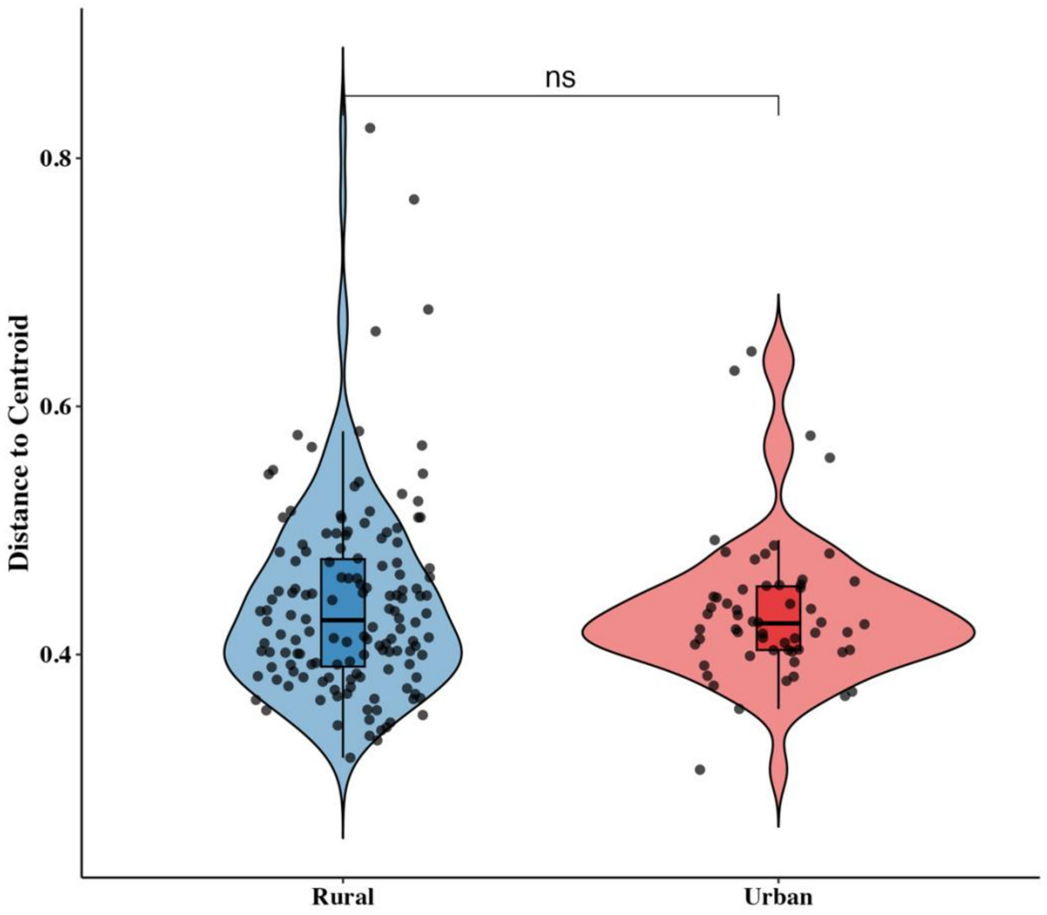


Supplementary Figure 3: Violin plot depicting the beta dispersion (distance to centroid) among the rural and urban dwellers.

Supplementary Tables legends

Supplementary Table 1: Demographic information of the volunteers
Supplementary Table 2: Differentially abundant ASVs among rural and urban dwellers
Supplementary Table 3: Common ASVs in rice beer and gut samples
Supplementary Table 4 : Differentially abundant functional pathways
Supplementary Table 5: BioSample IDs of the sequences
